# Assessment of the use and quick preparation of saliva for rapid microbiological diagnosis of COVID-19

**DOI:** 10.1101/2020.06.25.172734

**Authors:** Mariana Fernández-Pittol, Juan Carlos Hurtado, Estela Moreno-García, Elisa Rubio, Mireia Navarro, Marta Valiente, Aida Peiró, Alicia Capón, Nuria Seijas, Miguel J. Martínez, Climent Casals-Pascual, Jordi Vila

## Abstract

The objective of this study was to assess the performance of direct real time RT-PCR detection of SARS-CoV-2 in heated saliva samples, avoiding the RNA isolation step. Oropharyngeal and nasopharyngeal swabs together with saliva samples were obtained from 51 patients clinically diagnosed as potentially having COVID-19. Two different methods were compared: 1. RNA was extracted from 500 μl of sample using a MagNA Pure Compact Instrument with an elution volume of 50μl and 2. 700µL of saliva were heat-inactivated at 96°C for 15 minutes, and directly subjected to RT-PCR. One step real time RT-PCR was performed using 5 μl of extracted RNA or directly from 5 μl of heated sample. RT-PCR was performed targeting the SARS-CoV-2 envelope (E) gene region. Diagnostic performance was assessed using the results of the RT-PCR from nasopharyngeal and oropharyngeal swabs as the gold standard. The overall sensitivity, specificity, positive and negative predictive values were 81.08%, 92.86%, 96.77% and 65.00%, respectively when RNA extraction was included in the protocol with saliva, whereas sensitivity, specificity, positive and negative predictive values were 83.78%, 92.86%, 68.42% and 96.88%, respectively, for the heat-inactivation protocol. However, when the analysis was performed exclusively on saliva samples with a limited time from the onset of symptoms (<9 days, N=28), these values were 90%, 87.5%, 44% and 98.75% for the heat-inactivation protocol. The study showed that RT-PCR can be performed using saliva in an RNA extraction free protocol, showing good sensitivity and specificity.

## Introduction

COVID-19 is a very devastating pandemic infection caused by SARS-CoV-2, which originated in China and has currently spread all over the world^1^. Rapid diagnosis of COVID-19 is essential for the management of patients mainly in the emergency department^2^. Although some assays based on antigen-antibody reaction have been commercialized to detect either antigens or antibodies (IgA, IgM or IgG), the most sensitive tool to detect SARS-CoV-2 continues to be reverse-transcription real time polymerase chain reaction (RT-PCR) which specifically amplifies different genes encoded in the viral RNA from nasopharyngeal and/oropharyngeal swabs.

Shortages in swabs for collecting nasopharyngeal and oropharyngeal samples as well as insufficient RNA extraction kits can lead to a critical situation during a pandemic in which a huge number of samples must be processed. Therefore, the main objective of this study was to evaluate the use of saliva as an alternative and easier-to-collect clinical sample to detect COVID-19. The diagnostic performance of nasopharyngeal and oropharyngeal swabs was compared to the performance of direct heated saliva to RNA extracted samples.

## Materials and methods

### Patients

Consecutive patients attending the Emergency Department at the Hospital Clinic of Barcelona with laboratory or clinical-radiologic findings compatible with a diagnosis of COVID-19 were included in the study. Patients presenting infections or lesions in the oropharyngeal area were excluded. All patients included in the study had a primary diagnosis of COVID-19 by a RT-PCR from oropharyngeal and nasopharyngeal swabs samples. Oropharyngeal and nasopharyngeal swabs together with saliva samples were obtained from 51 patients clinically diagnosed as potentially having COVID-19.

### Samples and procedure

Nasopharyngeal and oropharyngeal swabs were deposited in a tube with 2 ml lysis buffer (guanidine thiocyanate, 2M; sodium citrate pH 7.0, 30 mM; dithiothreitol, 2 mM and triton X-100, 1%). All patients were asked to provide a saliva sample from the posterior oropharynx before tooth brushing and meal intake. Patients were instructed and supervised by a medical care team. Saliva samples with a volume less than 500μL were not included in the study. The samples were transported to the Clinical Microbiology Department of the Hospital Clinic in Barcelona, Spain, within less than 2 hours after collection. All saliva samples were stored at −80°C and processed together.

Two different methods were compared. In the first method, 350 µL of saliva were mixed with 350µL of lysis buffer (MagNA Pure Compact RNA Isolation Kit, Roche). RNA was extracted from 500 μl of sample using a MagNA Pure Compact Instrument (Roche, Basel, Switzerland) with an elution volume of 50μl. In the second method, 700µL of saliva were heat-inactivated at 96°C for 15 minutes, and 5 μl were subjected directly to RT-PCR. One step real time RT-PCR was performed using the RNA Process Control Kit (Roche, Basel, Switzerland) with 5 μl of extracted RNA or directly from 5 μl of heated sample. RT-PCR was performed targeting the SARS-CoV-2 envelope (E) gene region^3^. Diagnostic performance was assessed using the results of the RT-PCR from nasopharyngeal and oropharyngeal swabs as the gold standard.

### Statistical analysis

Statistical analyses were conducted using Stata 16.0 (*lroc* and *lstat* functions) and positive and negative-predictive (post-test) values were calculated for a disease prevalence of 10%.

## Results

A total of 51 patients with suspicious of COVID-19 were included in the study (Supplementary data 1), 37 patients gave positive by RT-PCR using oro- and naso-pharyngeal swabs, whereas 14 were negative. The overall sensitivity, specificity, positive and negative predictive values were 81.08%, 92.86%, 96.77% and 65.00%, respectively when RNA extraction was included in the protocol with saliva, whereas sensitivity, specificity, positive and negative predictive values were 83.78%, 92.86%, 68.42% and 96.88%, respectively, for the heat-inactivation protocol (Supplementary data 2). However, when the analysis was performed exclusively on saliva samples (heated only) with a limited time from the onset of symptoms (<9 days, N=28), these values were 90%, 87.5%, 44% and 98.75% for the heat-inactivation protocol (Table).

## Discussion

Two studies have recently evaluated the use of saliva for diagnosing SARS-CoV-2 infection, showing that the saliva viral load was highest during the first week after symptom onset and subsequently declined over time, which suggests that saliva may be a good non-invasive sample for detecting the presence of SARS-CoV-2 during the first days after the onset of symptoms^4,5^. Using RT-PCR, Pasomsub*et al*.^6^ found a sensitivity and specificity for saliva samples of 84.2% [95% confidence interval (CI) 60.4%-96.6%], and 98.9% (95% CI 96.1%-99.9%), respectively. Analysis of the two specimens demonstrated 97.5% of agreement (kappa coefficient 0.851, 95% CI 0.723-0.979; p <0.001). Moreover, it has also been suggested that saliva could be a more sensitive alternative to nasopharyngeal swabs^7^. Our results are in agreement with the abovementioned studies, but in addition we show that RT-PCR can be performed using an RNA extraction-free protocol with 91.9% of concordance with oropharyngeal and nasopharyngeal swabs (considering only samples collected below 9 days of the onset of the symptoms) This protocol modification reduced the turnaround time by 40 minutes, taking into account the 10 minutes of pre-processing plus 30 minutes of RNA extraction. In addition, the cost is also decreased. Moreover, the combination of a test with a high negative predictive value, and the simplified logistics of sample collection (patients provide the saliva samples with no need for personal protective equipment) is especially helpful to rule out infection during times of low incidence and also in low-resource settings. Larger studies are needed to prospectively validate these findings, and standardized saliva sample collection protocols are also necessary prior to implementation in the clinical setting.

---

**Table.** Sensitivity and specificity of RT-PCR in saliva sample using heat-inactivation protocol

| Saliva (Heating only) | % (95%CI) | % (95%CI) |
| --- | --- | --- |
|  | all days (N=51) | <9 days (N=28) |
| Sensitivity | 83.7 (67.9-93.8) | 90.0 (68.3-98.7) |
| Specificity | 92.8 (66.1-99.8) | 87.5 (47.3-99.6) |
| Positive likelihood ratio | 11.7 (1.76-77.9) | 7.2 (1.14-45.3) |
| Negative likelihood ratio | 0.17 (0.08-0.37) | 0.11 (0.03-0.44) |
| Accuracy | 91.9 (80.8-97.7) | 87.7 (69.8-97) |

